# Unsupervised functional neurocartography of the *Aplysia* buccal ganglion

**DOI:** 10.1101/2021.12.20.473553

**Authors:** Renan M. Costa, Vijay A. Dharmaraj, Ryota Homma, Curtis L. Neveu, William B. Kristan, John H. Byrne

**Author notes:** These authors contributed equally to the work (Renan M. Costa, Vijay A. Dharmaraj, Ryota Homma). **Correspondence should be addressed to:** John H. Byrne, Department of Neurobiology and Anatomy, McGovern Medical School at The University of Texas Health Science Center at Houston, 6431 Fannin Street, Suite MSB 7.046, Houston, TX 77030. **Author contributions:** Conceptualization: RMC, VAD, RH, CN, JHB. Methodology: RMC, CN, JHB. Investigation: RMC. Data curation: RMC, VAD. Software: RMC, VAD, RH. Visualization: RMC, RH. Analysis: RMC, VAD, RH. Funding acquisition: JHB. Project administration: JHB. Supervision: JHB. Writing—original draft: RMC, VAD, RH. Writing— review & editing: RMC, RH, CN, WBK, JHB. Michael E. DeBakey Department of Surgery, Baylor College of Medicine; Houston, United States.

## Abstract

A major limitation of large-scale neuronal recordings is the difficulty in locating homologous neurons in different subjects, referred to as the “correspondence” issue. This issue stems, at least in part, from the lack of a unique feature that unequivocally identifies each neuron. One promising approach to this problem is the functional neurocartography framework developed by Frady et al. (2016), in which neurons are identified by a semisupervised machine learning algorithm using a combination of multiple selected features. Here, the framework was adapted to the buccal ganglia of *Aplysia*. Multiple features were derived from neuronal activity during motor pattern generation, responses to peripheral nerve stimulation, and the spatial properties of each cell. The feature set was optimized based on its potential usefulness in discriminating neurons from each other, and then used to match putatively homologous neurons across subjects with the functional neurocartography software. An alternative matching method based on a cyclic matching algorithm was also developed, which allows for unsupervised extraction of groups of neurons and automated selection of high-quality matches. This improvement enabled unsupervised implementation of the machine learning algorithm, thereby enhancing scalability of the analysis. This study paves the way for investigating the roles of both well-characterized and previously uncharacterized neurons in *Aplysia*, and helps to adapt the neurocartography framework to other systems.

## INTRODUCTION

Analyses of the function and connectivity of a neuronal circuit require the ability to identify and characterize the individual functional units within it. This characterization is facilitated in simpler nervous systems because they are known to contain many individually unique neurons (e.g., Byrne, 2019) that are regarded as functional units. Conversely, in more complex systems, the functional unit of interest may be a group of similarly-behaving neurons or even an entire region. In either case, the traditional approach has been to distinguish these functional units by using some defining feature or trait (e.g., a protein, gene, metabolite, neurotransmitter, morphology, synaptic connection, or activity pattern). However, in the majority of cases neurons or groups of neurons are not readily identifiable by a signature feature alone. Even when a single feature purportedly identifies a neuron type, future experiments tend to find subcategories of that group (e.g., Abrahao and Lovinger, 2018; Björklund and Dunnett, 2007; Hernández et al., 2015; Mallet et al., 2012; Tiklová et al., 2019). A notable example of the problem is comparing the same circuit between two different subjects. Although investigation *in vitro* may demonstrate two subjects to be physiologically similar, inherent intersubject variability can make identifying even the most obvious homologous functional units between them nontrivial. The problem of identifying homologous units in spite of intersubject variability is known as the issue of correspondence (Aberg et al., 2009; Hassanien et al., 2009; Yendiki et al., 2016; Yu et al., 2021).

To address the issue of correspondence, Frady et al. (2016) developed an analytical framework termed *functional neurocartography*. They built and optimized a rich feature-space composed of several features that were informative for the identification of many individual leech neurons recorded using voltage-sensitive dye (VSD) imaging. Cells were algorithmically compared based on features extracted from neuronal activity during behavior, and putatively homologous neurons were identified through careful curation of the results. Neurons from these experiments were then fit onto a reference map template based on known information from the literature and this mapping process also revealed previously uncharacterized neurons. This approach is effective, but requires the curator to be well informed about the structure and function of the neuronal circuit being investigated. The present study aimed to generalize neurocartography to nervous systems where such information may not be readily available.

This study applied the functional neurocartography approach to the neurons in the buccal ganglion of the marine mollusc *Aplysia californica*. The buccal ganglia contain a central pattern generating (CPG) circuit that mediates the consummatory aspects of feeding behavior. The ganglion continues to produce spontaneous buccal motor patterns (BMPs) after removal from the animal, and many neurons essential for the generation of BMPs have been characterized (for reviews, see Baxter and Byrne, 2006; Cropper et al., 2004). In addition, VSD imaging has been successfully used in large-scale recordings from buccal ganglia neurons (Costa et al., 2022; Morton et al., 1991; Neveu et al., 2017). Although neurons in the buccal ganglion are individually identifiable, the criteria vary considerably. For the majority of neurons, an elaborate combination of anatomical and physiological properties are necessary for identification. Many neurons remain uncharacterized because their identifying features have yet to be documented. Searches for uncharacterized neurons that contribute to BMPs are still ongoing (e.g., Costa et al., 2022; Momohara et al., 2022). Consequently, this model system could benefit greatly from functional neurocartography.

Here, we implemented and expanded the functional neurocartography framework, that is, obtained anatomical and physiological neuronal features that were tailored to the *Aplysia* buccal ganglion and identified matched neurons among preparations based on those features. Several key adaptations were made. First, in addition to the typical VSD imaging of neuronal activity during motor pattern generation, VSD signals were used to measure neuronal responses to stimulation of different peripheral nerves. This manipulation added a set of robust and distinct features to the feature-space. Second, a novel tool was developed for matching neurons across subjects. A cyclic matching algorithm (Hofbauer, 2016) was used to chain together pairs of experiments and form families of neurons. This approach circumvents the problem of integrating pairwise matching results across all animals (termed “stitching” in the functional neurocartography framework) and enables an unsupervised analytical pipeline for the cross-subject matching of homologous neurons. This automated analytical pipeline helps enhance the scalability of the framework to accommodate a larger number of subjects. These adaptations and enhancements will facilitate the future use of neurocartography for analyses of the *Aplysia* buccal ganglion, as well as other nervous systems.

## RESULTS

### Spatial and functional features

In order to build an information-rich feature-space, we first collected data, defined many spatial and functional features of each neuron, and assessed the usefulness of each feature. Potential sources of useful information on individual neurons include size and position, characteristics of activation during motor pattern generation, and responses to stimulation of peripheral nerves. This information was acquired by combining population-wide VSD imaging with extracellular nerve recording and stimulation. Here, we provide an overview of the experimental configuration and of the extracted neuronal features. Detailed descriptions are provided in Materials and Methods.

The activity of individual neurons in isolated *Aplysia* buccal ganglia during motor pattern generation (Fig. 1) was measured by staining ganglia with VSD RH-155. Two sets of experiments were performed: one recording from the caudal (n = 7) surface of the left buccal hemiganglion and the other from the rostral (n = 6) surface. Suction electrodes were used to record activity in buccal nerves 1–3 (cBn1, cBn2, cBn3, iBn2, iBn3—where *c* and *i* denote whether the nerve was *contralateral* or *ipsilateral* to the imaged hemiganglion), the esophageal nerve 2 (iEn2) and the radular nerve (iRn). Activity in these nerves is correlated with specific phases of the animal’s feeding behavior (i.e., protraction, retraction, post-retraction; Cropper et al., 2004; Morton and Chiel, 1993b). The relationship between the activity of individual neurons and these phases, in turn, provides potentially useful information about the behavioral role of each cell (e.g., Morton and Chiel, 1993a).

**Figure 1.**
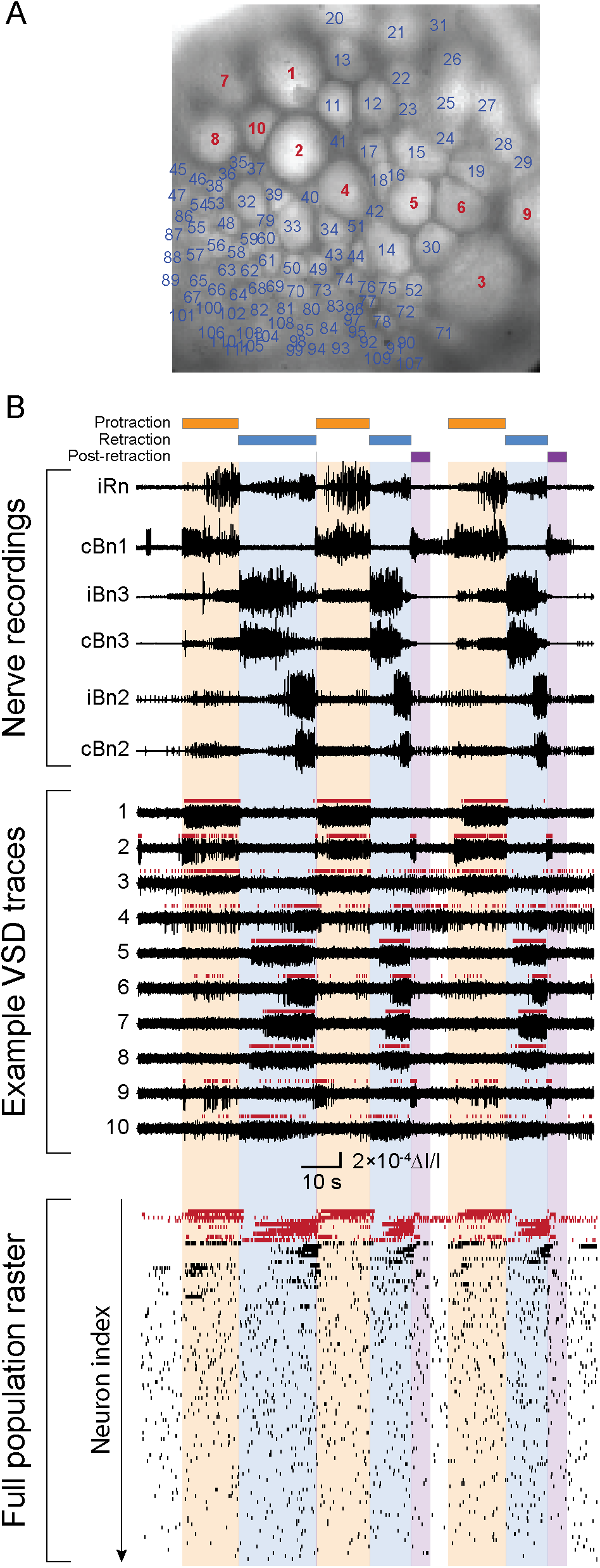
Concurrent voltage-sensitive dye (VSD) imaging and extracellular nerve recordings during motor pattern generation. **A.** Caudal surface of the buccal ganglion. Example neurons displayed in (B) are highlighted in red. A total of 111 neurons were in the field of view for this preparation. **B.** (Top) Extracellular nerve recordings were used to monitor motor pattern generation and to identify each phase of a motor pattern (protraction, retraction and post-retraction; respectively orange, blue and purple bars and shading). (Center) Example VSD traces from 10 neurons during the same period. Vertical red lines above each trace denote the occurrence of action potentials. (Bottom) Raster of the spike trains from all 111 neurons. Neurons displayed as examples above are highlighted in red.

The timing of the spike activity detected from the VSD recordings was examined (Fig. 1B; see Material and Methods). Many neurons were active predominantly during one or more of the three BMP phases (protraction, retraction, post-retraction). Phase preference was quantified as the normalized activity of a neuron during each phase of the BMP (Fig. 2A). This yielded three *preference* features—one for each BMP phase.

**Figure 2.**
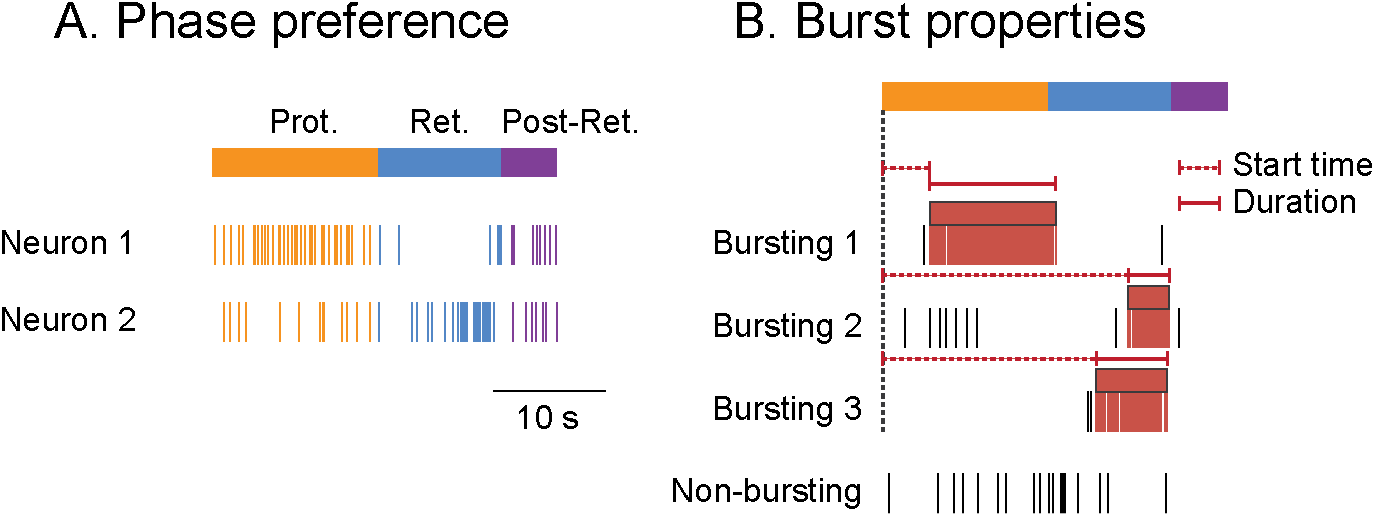
Features of neuronal activity during motor patterns. **A.** *Preference* of neuron activity for each phase of a motor pattern. Orange box denotes protraction, blue retraction, and purple post-retraction. Spikes that were used to compute a neuron’s preference toward a given phase are highlighted in the corresponding color. **B.** Bursts of activity in each spike train were labeled and neurons were classified as either bursting or non-bursting. Red boxes denote bursts identified in each spike train. Spikes in each burst are highlighted in red. The overall level of burst-like activity in each neuron was quantified as the firing rate during bursts. This feature was termed *frequency. Start time* was the time elapsed between the onset of a motor pattern and the onset of burst-like activity in each neuron. *Duration* was the total length of bursts for each neuron. See Materials and Methods for details on the quantification of each feature.

Neurons were classified as bursting or non-bursting using previously described criteria (Neveu et al., 2017; see Materials and Methods), which yielded a binary feature termed *bursting*. Additional features were obtained from the burst properties of each *bursting*-positive neuron, including the firing rate during bursts (*frequency*), and the *start time* and *duration* of burst-like activity during BMPs (Fig. 2B). *Start time* and *duration* were extracted after normalizing the duration of each phase of a motor pattern, whereas *frequency* was calculated prior to normalization.

Neurons within the buccal ganglia may send processes through one or more peripheral nerves (Evans and Cropper, 1998; McManus et al., 2014; Morton and Chiel, 1993b; Scott et al., 1991), as well as receive synaptic input from cells whose processes pass through these nerves (Nargeot et al., 1999). Nerve stimulation can therefore activate neurons both directly and indirectly, thereby providing a potential rich source of information on many cells. VSD imaging was used to capture the responses of all neurons to stimulation of each of the recorded nerves (Fig. 3). The amplitude of neuronal activity following nerve stimulation yielded seven *nerve response* features, one for each of the seven stimulated nerves (cBn1–3, iBn2–3, iEn2, iRn).

**Figure 3.**
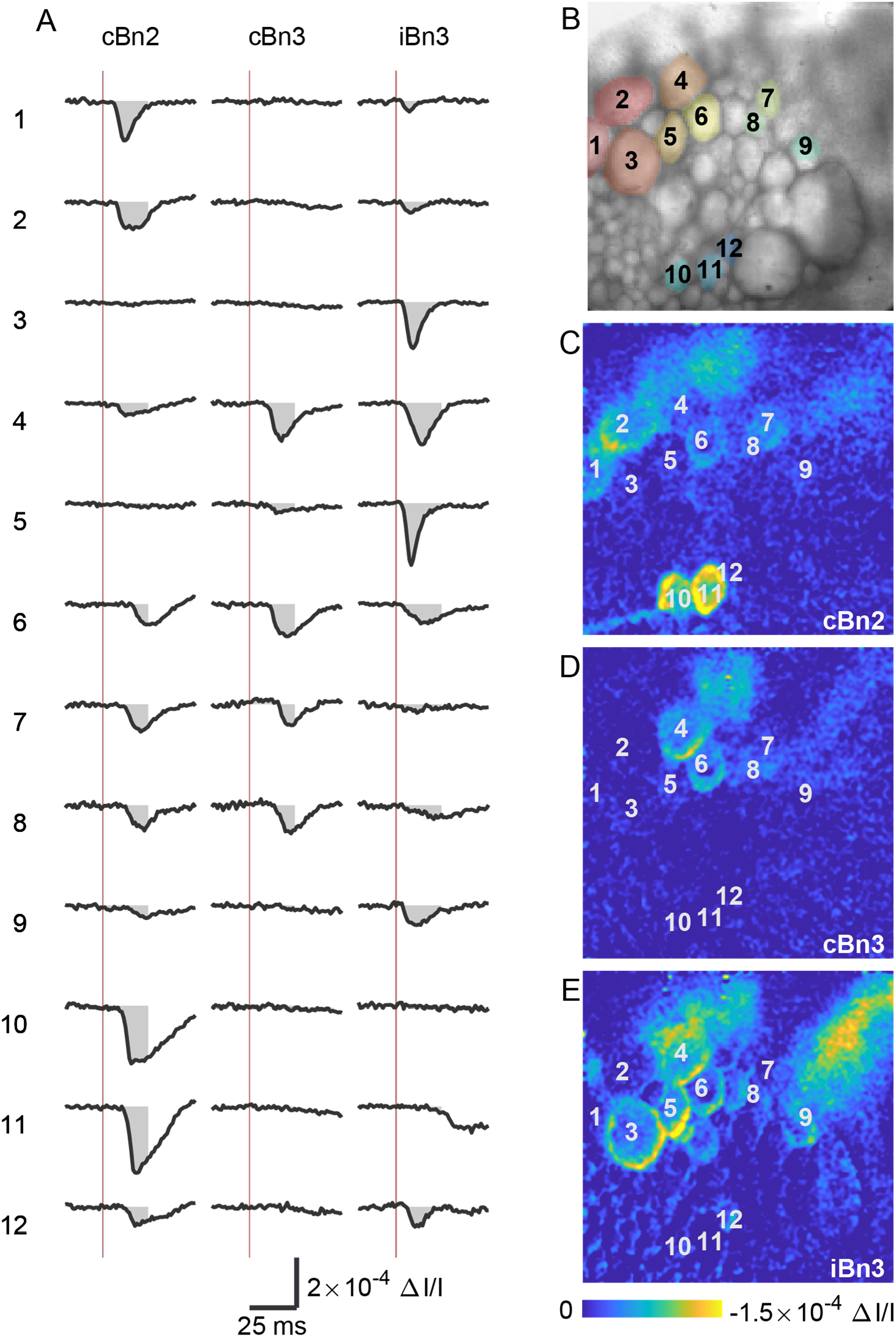
Feature extraction from responses to nerve stimulation. Following stimulation of individual nerves, distinct subsets of neurons exhibit VSD responses. The *response amplitude* feature was defined as the integral of the -ΔI/I trace over the first 25 ms following nerve stimulation (shaded area in A). **A.** Example time courses of the VSD signal in response to the nerve stimulations. Note that a downward deflection in the VSD trace corresponds to a depolarization. Responses of 12 neurons to 3 nerves (cBn2, cBn3, and iBn3) are presented. Neuron indexes correspond to the numbers in panels B–E. Each trace is an average of 40 responses. Red vertical lines indicate the time of nerve stimulation. **B.** Image of the VSD-stained buccal ganglion showing the 12 neurons that generated the recordings in panel A. **C–E.** Spatial patterns of activity in response to stimulation of each of the three nerves. The response maps were obtained by subtracting the ΔI/I image of the baseline periods (average over −10–0 ms and 45–55 ms relative to the stimulus onset) from that of the post-stimulus period (average over 45 ms after the stimulus; a larger time window than the 25 ms window for the amplitude parameter was used for improved image quality).

Finally, the spatial features for all imaged neurons were parametrized. The ganglion was positioned at approximately the same location across experiments. In addition, variations were accounted for by approximating the orientation and offset of both the ventral cluster and the entire ganglion, and then aligning each recording accordingly, as previously described (Neveu et al., 2017). The centroid of the pixels overlying each neuron was used as the *x* and *y* coordinates for the *position* feature, and the total number of pixels was used as the *size* feature.

### Feature assessment

We next evaluated what subset of features would form the optimal feature-space for the purpose of identifying putatively homologous neurons. The quality of a feature depends primarily on three factors: i) how much information it provides; ii) how much of that information is unique relative to other features (non-redundancy); and iii) the extent of its variability within and across subjects (consistency). Thus, a high-quality feature must offer non-redundant information that is sufficiently consistent to allow neurons to be distinguished from one another.

The quality of the feature-space was quantified using a metric (termed *discriminability index*) that reflects the percentage of neurons that are distinguishable in a certain feature-space (Frady et al., 2016). Briefly, this index asks whether, upon splitting the data in two halves, a given neuron is closer to itself than to other similar neurons in the feature-space. If that is the case, the neuron is deemed discriminable, that is, distinguishable from other neurons. The discriminability index was computed for systematically manipulated possible subsets of features (Fig. 4; see Materials and Methods). By selecting the subset of features that maximizes the number of discriminable neurons, we attempted to optimize the usefulness of our feature-space.

**Figure 4.**
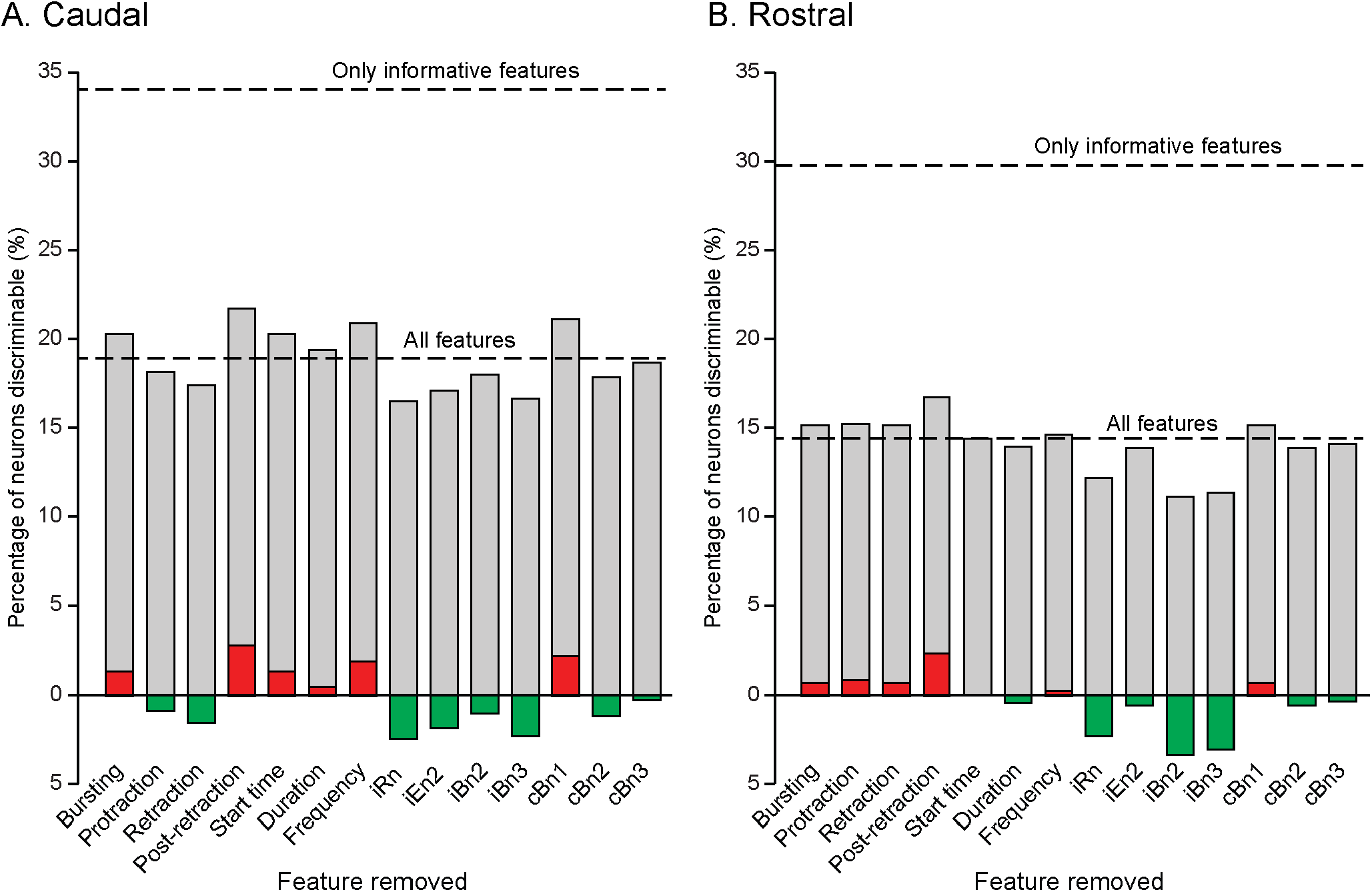
Effect of individual feature removal on neuron discriminability. **A.** Discriminability of neurons visible from the caudal surface. If a feature contains useful information, removing it must necessarily reduce the ability to discriminate neurons within the subject (green). On the other hand, if removing a feature improves (red) discriminability, it should not be included in the feature-space. Dashed lines represent the discriminability of feature-spaces containing either all features, or only those deemed informative. **B.** Same as (A), but for neurons visible from the rostral surface.

Feature assessment was performed separately for the rostral and caudal buccal neurons, and resulted in two distinct subsets of features. For caudal preparations, the selected feature-space consisted of protraction and retraction phase preferences, the start time burst property, and nerve responses for iRn, iBn2, iBn3, cBn2, and cBn3. Conversely, the feature-space for rostral preparations consisted of start time and duration burst properties, in addition to nerve responses for iRn, iBn2, iBn3, cBn2, and cBn3. Responses to iEn2 stimulation were small compared to other nerves, and we could not rule out the possibility that these responses were due to noise. Therefore, we conservatively chose not to include responses to iEn2 stimulation in the final feature-spaces, even though this feature was classified as useful for both caudal and rostral populations by the discriminability analysis, and thus potentially contained valuable information. By default, both caudal and rostral spaces included size and position features. The differences between the feature-spaces yielding highest discriminability for each surface imply that certain information may be of greater or lesser value depending on the neuronal population of interest. Specifically, the rostral neurons appeared to rely more on the temporal properties of bursts, while caudal neurons relied more on BMP phase preference. Both, however, were equally reliant on the information contained within the nerve response features. The differences in the optimal subsets of features further substantiated the use of the discriminability index for feature quality assessment and, in addition, highlighted the underlying physiological differences between the rostral and caudal sides of the buccal ganglia.

### Pairwise functional neurocartography

The assessment of feature discriminability yielded a caudal and a rostral libraries of neurons with respective feature sets selected for intersubject comparison. Pairwise comparisons of neurons were performed between every unique pair of experiments for each of the caudal and rostral datasets as the first step for identifying homologous neurons across experiments. These pairwise comparisons, performed with the Imaging Computational Microscope (ICM) module of the neurocartography toolset (Frady and Kristan, 2015), generated a metric of neuron-to-neuron match strength, which was based on the extent of similarity across all input features as well as quantity and quality of alternative candidates for the match. The ICM then uses the Hungarian algorithm (Kuhn, 1955) to ensure that as many neurons as possible find matches between experiments by allowing non-best matches. The software is also capable of optimizing the relative contributions of each feature (feature skewing) to manage intersubject variability. Feature skewing was not utilized at this stage, but was used in subsequent analyses (see the following subsections).

One drawback of the pairwise approach is that information across all possible pairs needs to be “stitched” (i.e., integrated in some way to obtain unified insights; Frady et al., 2016). Stitching is nontrivial because full consistency of pairwise matches among three or more animals is not guaranteed (Fig. 5A). Frady et al. addressed this issue by training the matching and feature skewing algorithm with human instructions, and then manually curating the results to identify “canonical” neurons that were consistently matched across animals. Although this approach offers the highest possible accuracy, it relies on expertise, and requires substantial human effort. The latter is a limiting factor for the number of animals to which the approach can be applied, as the number of possible pairs of animals is equal to N×(N-1)/2 (i.e., N choose 2), where the N is number of animals. Thus, supervision effort (i.e., the number of matches the analyst has to judge) scales rapidly with the number of animals.

**Figure 5.**
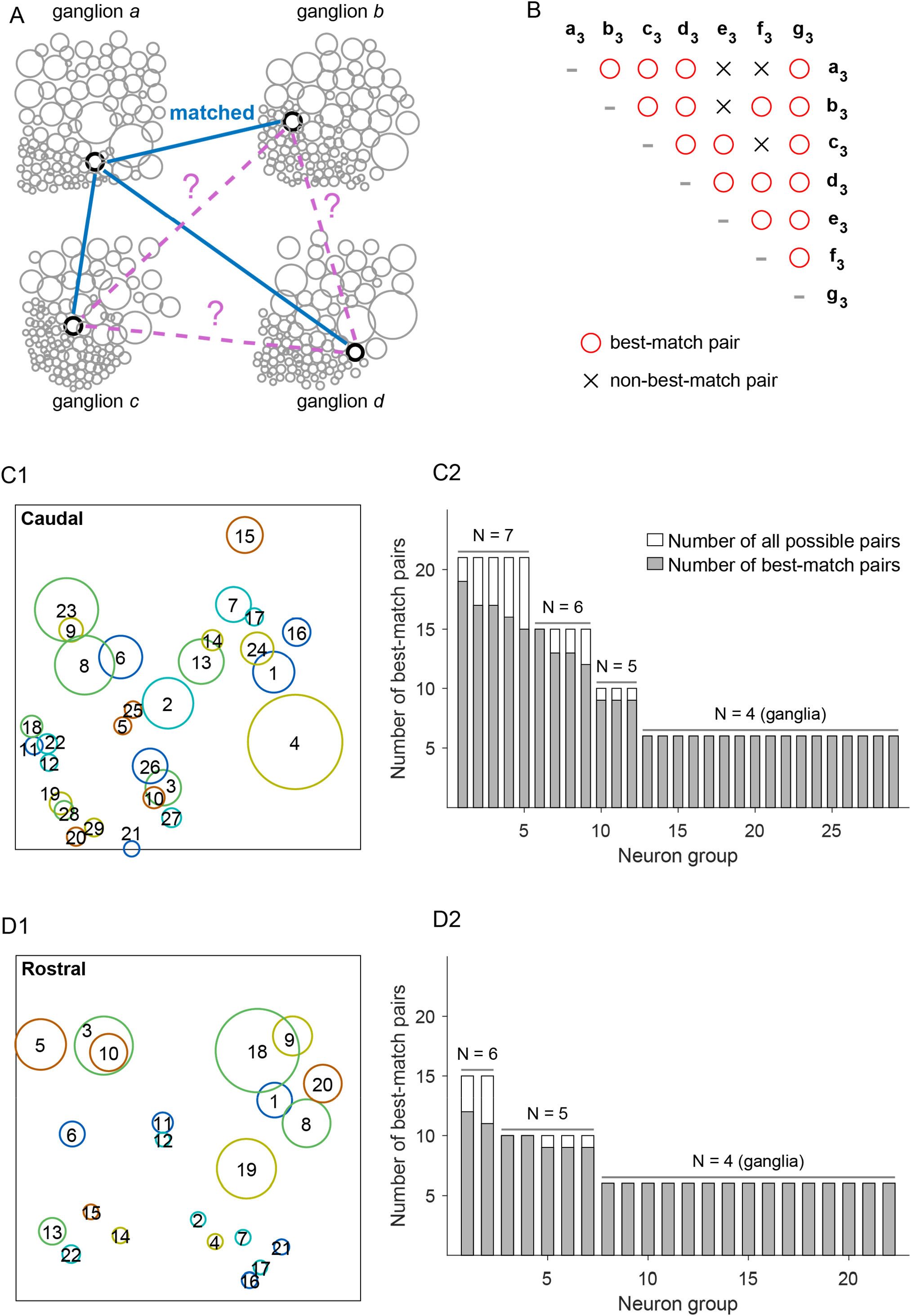
Pairwise matching analysis. The Imaging Computational Microscope (ICM; Frady and Kristan, 2015) was used for matching neurons between every pair of experiments based on neurons’ similarities in their spatial and functional features. **A.** Matching neurons across ganglia. Although a given neuron in ganglion *a* will be matched with one neuron in each of ganglia *b*, *c*, and *d*, it is not guaranteed that these neurons in *b*, *c*, and *d* will, in turn, be each other’s best possible matches when their ganglia are directly compared. As a solution to this problem, neurons were considered to be in a matching group when all neurons were the best match (open circle, B) for each other in all possible pairwise comparisons (“perfect group”) among at least four ganglia. **B.** Groups with overlapping neurons were then merged into a single group. The diagram presents such an instance, where letters a_3_-g_3_ each indicate one neuron in each of the seven separate ganglia. (Specifically, neurons that fell into the group 3 in panel C.) For example, the four neurons a_3_, b_3_, c_3_, and d_3_ form best-match pairs (red circles) a_3_-b_3_, a_3_-c_3_, a_3_-d_3_, b_3_-c_3_, b_3_-d_3_, and c_3_-d_3_. The seven neurons contained eight of such perfect groups (abcd, abcg, abdg, acdg, bcdg, bdfg, cdeg, and defg; subscripts are omitted for clarity) and were merged into a single group. Note that several pairs in the resulting single group are non-best-matches in direct pairwise comparisons (gray crosses). In this example, out of 21 possible pairs between 7 neurons, 17 pairs are best-matches and 4 are non-best. **C1.** All caudal-side groups of neurons identified through the aforementioned steps. Each circle represents a single group, and the position and size of circle show the median position and size of participating neurons. The colors of circles are for visibility and do not represent specific information. **C2.** The number of participating neurons and number of best-match pairs among them are presented for every group in C1. Each bar represents a single group and its position along the X-axis corresponds to the group number in C1. **D.** The identified groups from six rostral-side experiments presented in the same format as in C. These results are based on the unskewed feature-space (see Figs. 7 and 8).

Consequently, a more automatic approach for the stitching process was sought. Here, matches were assessed based solely on the consistency of pairwise matching across animals. A set of four neurons was considered successfully matched across subjects when every possible pairing among them was the best possible match for each neuron in pairwise matching (termed “perfect group”). Four was chosen as the minimum majority of 7 caudal experiments or 6 rostral experiments so that the corresponding neurons cannot appear in two separate sub-groups. Furthermore, when two or more such groups of four neurons shared at least one member neuron, these groups were merged even if there were pairs of non-best match neurons among the groups (Fig. 5B). Before the optimization of intersubject variability, this procedure identified 29 groups in the caudal dataset (Fig. 5C, D) and 22 groups in the rostral dataset (Fig. 5E, F). These numbers are not distant from what is to be expected from the discriminability analysis above, although they are not intended for direct comparison. For the caudal and rostral datasets, 12/29 and 7/22 groups, respectively, contained five or more member neurons after group merging.

### Cyclic matching and copresence analysis

The approach above allows integration of pairwise matches based on the cross-animal consistency of identified groups of neurons, with minimal human effort. However, this solution does not provide an estimate of error, and thus contains little information about the level of confidence in individual identified groups. For example, some of the groups (in particular, those with four members) may be false positives that were matched by chance. In addition, the criterion of perfect groups is possibly too stringent when the number of subjects (and thus its minimum majority) becomes larger, implying another challenge for a scalable automated analysis.

To overcome this limitation, we applied a fundamentally different matching strategy, termed *d*-dimensional stable matching with cyclic preferences (cyclic matching; Hofbauer, 2016), in which matching of more than two members can be achieved at once by considering the preference of members in a unidirectional and cyclic manner (Fig. 6A; see Materials and Methods). This cyclic matching algorithm yields complete sets of neurons (referred to as *families*) in the sense that each set contains neurons from all subjects. However, the matching results generally depend on the unidirectional order of experiments chosen to compute cyclic preferences. Thus, cyclic matching was performed repeatedly using randomly shuffled orders of experiments, and the robustness of resulting matches was assessed based on their consistency across runs (Fig. 6B). Specifically, the number of times a pair of neurons was present in the same family out of 100 repetitions was counted (referred to as *copresence count*). The copresence counts collectively form a set of copresence profiles for each reference neuron (Fig. 6B). When a specific neuron exhibits a high copresence count with respect to the reference neuron, they are considered a strong match. When two neurons in the same subject show comparable copresence counts, or when none of the neurons have a salient copresence count, the matching counterpart is uncertain. By taking every neuron in the dataset as the reference neuron, putatively matching pairs of neurons were identified and a confidence metric was assigned to each match that could be used in subsequent analyses (i.e., the copresence count).

**Figure 6.**
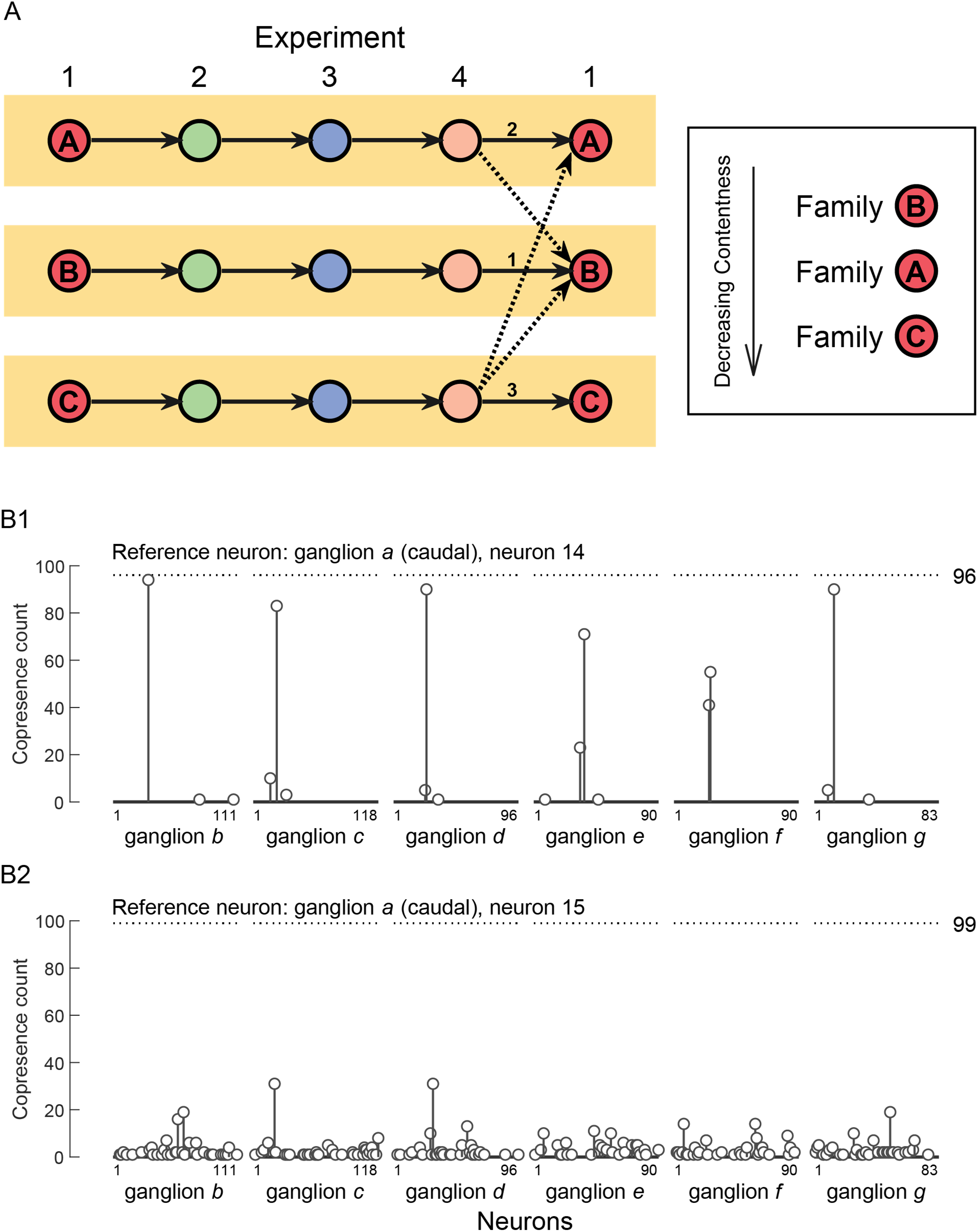
Cyclic matching and copresence analysis. **A.** Schematic diagram presenting the concept of the cyclic matching algorithm. It relies on sequential pairwise matches generated by the Hungarian algorithm, and then assesses the preference of the last cell to the first cell of the same “family” as the rank of their similarity relative to the similarities between the last cell and each cell in the first experiment. The family with highest preference is removed from the dataset as an established family. The process is repeated until all cells (in the experiment with fewest cells) form a family. The order of experiments for matching can be arbitrary and generally matching results are not identical when the orders of experiments are different. **B.** The robustness of individual matchings was determined by performing the cyclic matching process many times (100 cycles) with randomly permuted orders of experiments. A specific neuron was picked as the reference and for every neuron from other experiments (i.e., neuron numbers in the x axis), the number of cycles in which both belonged to the same family was counted (“copresence”). Two examples of such an analysis are presented in two sets of stem plots. **B1,** the reference neuron was picked from ganglion *a* (see Fig. 5). There are six stem plots, each of which corresponds to one of ganglia *b–g*, and each stem represents a single neuron that has a non-zero copresence count. Although a single neuron had a high copresence in the vast majority of runs in ganglia *b*, *c*, *d*, and *g*, there were two competing neurons in ganglia *e* and *f*, suggesting a failure to find a reliable counterpart. **B2**, the reference was another neuron from ganglion *a*. In this example, there were many candidate neurons in all ganglia, implying that matching is unstable across runs. The same analysis was conducted for every neuron from every experiment, and a pair of neurons is considered as a reliable match when they reciprocally demonstrate highly exclusive copresence. The horizontal dashed line and number in the right indicate the number of cycles (out of 100) in which the reference neurons appeared in the matching results.

### Feature skewing for the optimization of intersubject comparisons

In our automation-oriented approaches of either pairwise matching or cyclic matching, one key component of the original neurocartography framework was left unused. The ICM software has the capability of optimizing intersubject comparisons through skewing of the feature-spaces of individual subjects. Feature-space skewing is designed to minimize the distances among a set of selected matches, which are deemed “correct.” In so doing, skewing emphasizes features that are informative for the correct matches and minimizes the impact of overall intersubject variability by de-emphasizing less informative features. But to implement skewing, it is necessary to have data on which matches should be selected as correct. Although such data was provided manually by the analyst in the original framework, the copresence results provide that instruction data in the present study. As the high copresence count implies highly consistent matching results across different runs of cyclic matching, it was considered as a proxy of the reliability and used to label the match as correct or not for this analysis.

Selected matches were determined using a copresence proportion criterion, computed over 100 repetitions of cyclic matching. For each match, the copresence proportion was the copresence count of the match (e.g., stems in Fig. 6B) divided by the appearances of the reference neuron (e.g., dashed lines in Fig. 6B). Matches that exceeded a given copresence proportion threshold were selected as “correct” for feature-space skewing. We compared various skewed and unskewed feature-spaces by using the sparseness of the full copresence profile of each reference neuron as an overall quality metric. Neurons whose matches are highly random across runs are expected to have low sparseness, whereas neurons that have consistent matches will have sparse copresence profiles. In other words, the sparseness is high when the copresence counts converge to one or just a few neurons, and is low when copresence counts distribute broadly among many neurons. Thus, if skewing results in copresence profiles that converge to fewer neurons, it increases the overall sparseness and improves the feature-space relative to intersubject variability. The ratio of the *ℓ*_2_ norm to the *ℓ*_1_ norm was chosen as the sparseness metric. Both norms are widely used to quantify sparseness, and the ratio between them is a convenient metric due to its scale-invariance (Krishnan et al., 2011; Yin et al., 2014).

A range of copresence proportion thresholds was explored. In caudal preparations, a threshold value of 0.3 yielded the highest overall sparseness (Fig. 7A), the highest number of cells showing a large change (>2.5 times the standard deviation) in sparseness (Fig. 7B) and led to smaller increases in sparseness in many other cells (Fig. 7C). The same threshold also had a similar performance on rostral preparations (Fig. 7D–E). Upon examining the copresence profiles of individual neurons before and after skewing (Fig. 7G–H), we found that many cells that were ambiguously matched across multiple runs in the unskewed feature-space converged more consistently onto the same matches after skewing. These results suggest that skewing to favor matches that are consistent across multiple cyclic runs can improve the feature-space. Thus, a copresence proportion threshold value of 0.3 was used for subsequent analyses of both caudal and rostral preparations.

**Figure 7.**
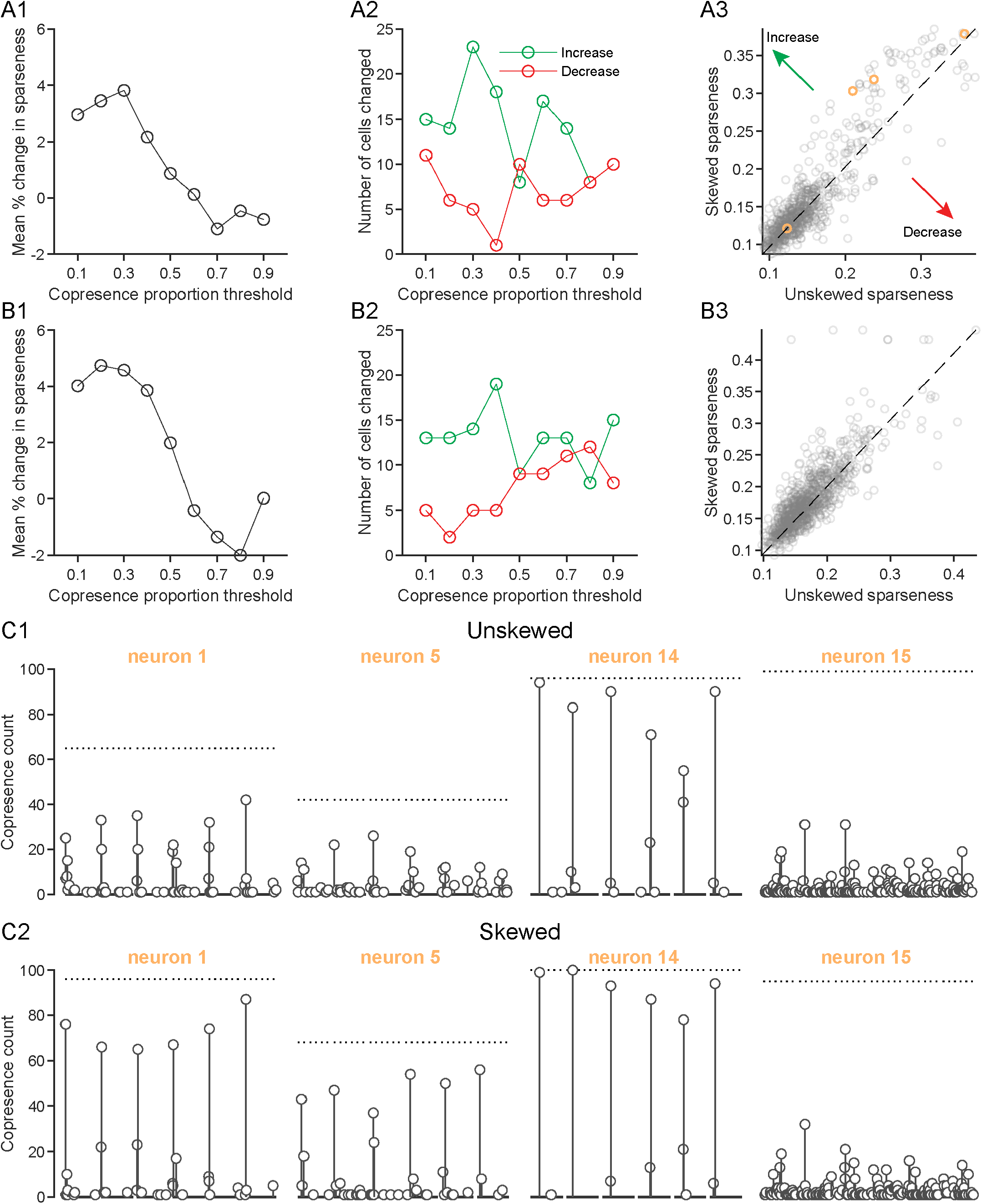
Feature-space skewing using the cyclic matching copresence criterion. A subset of cross-subject cell matches was automatically selected for skewing of the feature-space to optimize the overall cross-subject comparisons. Matches were included if they met a copresence criterion threshold. **A1.** Overall change in sparseness of the copresence profiles over a range of copresence criterion thresholds. Averaged across all cells from caudal preparations. The threshold value of 0.3 had the largest overall improvement in sparseness. **A2.** Total number of cells displaying a significant (>2.5 standard deviations) increase or decrease in sparseness for each threshold. **A3.** Sparseness of the copresence profile of individual neurons before and after skewing of the feature-space using the 0.3 threshold value. Neurons above the diagonal display increased sparseness after skewing, whereas those below display decreased sparseness. Cells highlighted in orange correspond to examples in panels C1 and C2. **B.** Same as A, but for rostral preparations. **C1.** Copresence profiles of four example reference neurons from caudal preparations with no skewing of the feature-space. Copresence profiles follow the same scheme and reference ganglion as Fig. 6. Note that plots are horizontally compressed to show all target ganglia for each neuron. **C2.** Copresence profiles of the same reference neurons after skewing of the feature-space using the 0.3 copresence criterion threshold.

### Groups of putatively homologous neurons based on cyclic matching and comparison with pairwise matching

We next compiled the results of cyclic matching to search for groups of putatively homologous neurons. To achieve this, we first extracted reliable pairs of matching neurons. Reliable pairs were determined based on the uniqueness (exclusiveness) of the copresence count in the copresence profile of a given experiment and a given reference neuron. We defined the uniqueness index (UI) (see Material and Methods), which takes values between 0 and 1 for neurons with highest copresence counts, and values between −1 and 0 for any other neuron. Higher values of UI indicate higher degrees of uniqueness. When a given neuron takes all copresence count in the experiment (i.e., completely exclusive), UI is equal to one, and when two neurons share the same highest count, their UIs are equal to zero, for example. A pair of neurons were considered as reliably matched when they reciprocally had a UI above 0.5 (see Material and Methods). Finally, reliable pairs that shared the same neurons were agglomerated to form a neuron group. In the cases in which the resulting groups included two neurons from the same subject, both neurons were eliminated from the group. Groups with four or more member neurons were kept for further analyses, although it was technically possible to identify groups of fewer member neurons.

The above process was applied to the cyclic matching results obtained after feature skewing. Nineteen groups from the caudal dataset (Fig. 8A–D), and 22 groups from the rostral dataset (Fig. 8E–F) were identified. For comparison, the cross-animal stitching of pairwise matches (i.e., Fig. 5C–D) was repeated using the feature-skewed datasets, which yielded 35 and 18 groups in caudal and rostral datasets, respectively.

**Figure 8.**
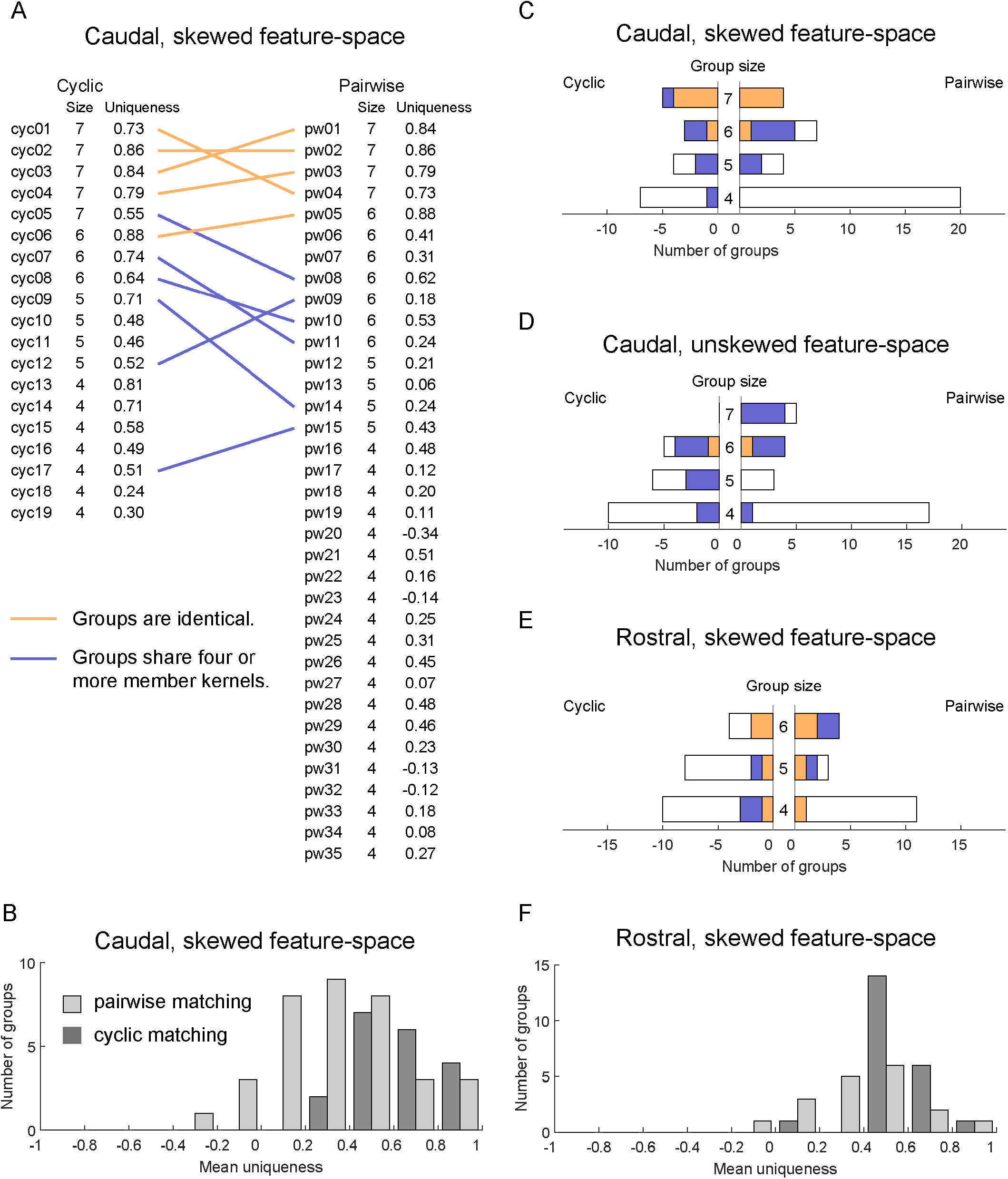
Assembly of the groups of putatively homologous neurons based on cyclic matching and optimized feature-space. By compiling the information of neuron pairs that showed highly exclusive copresence from the results of cyclic matching (see text), groups of putatively homologous neurons were identified. These groups were compared with the groups obtained through the cross-subject comparison of pairwise matching results. To ensure a meaningful comparison, pairwise matches were updated relative to those in Fig. 5 by using the optimized feature-space. **A.** Direct comparison of groups from cyclic matching and pairwise matching of the caudal dataset. Left and right rows list the neuron groups based on cyclic (cyc) matching and pairwise (pw) matching, respectively, with the number of member neurons and mean uniqueness index (a possible metric of matching quality). Groups connected with an orange line are identical in their member neurons. Groups connected with blue lines share four or more member neurons but also contain at least one inconsistent neuron in one or both of groups. **B.** Histograms of the distributions of the mean uniqueness indices of pairwise and cyclic groups. **C.** Summary of the number of established groups in panel A, as well as fully or partially consistent groups between cyclic and pairwise matching. **D.** Summary data of group numbers based on the unskewed feature set. Note that there is no group with all seven experiments in cyclic matching (left side), resulting in fewer fully consistent groups compared to C. **E, F.** Summary of group numbers and histograms of uniqueness indices in the rostral dataset. The format of graphs is the same as panels C and B, respectively.

To compare cyclic groups and pairwise groups, we counted groups that had identical member neurons, as well as those that were not identical, but still shared four or more members. In the caudal dataset, there were five identical groups and six groups with 4+ shared members (Fig. 8A, C). The same analysis was performed using the dataset without feature skewing and obtained one identical group and seven groups with 4+ shared members (Fig. 8D). Of note, no seven-member groups were obtained from cyclic matching with the unskewed data. The same comparison of group members was conducted in the rostral dataset (with feature-skewed data), which resulted in four identical groups and three groups with 4+ shared members (Fig. 8E). In addition, the uniqueness index of the group was calculated, which is defined as the mean uniqueness index calculated across all possible pairs (bidirectional) among member neurons in the group (Fig. 8A, B, F). Although the group uniqueness indexes remain relatively high in the cyclic matching groups, which was expected because the grouping process had set a threshold for uniqueness index, the indexes distributed more broadly in the pairwise matching groups. This difference might reflect the lack of a means for quality control in the cross-animal comparisons of pairwise matching.

Finally, the features of the putatively homologous neurons identified through cyclic matching for caudal and rostral preparations were examined. Homologous caudal neurons ranged from small to large, but most appeared to be on the larger end (Fig. 9A). Despite substantial variability in the exact position of individual neurons, several patterns can be noted. For example, neuron cyc05 (yellow, Fig. 9) was consistently located on the top left quadrant, often next to or near neurons cyc04 (magenta) and cyc08 (brown). Neurons such as cyc02 (green) and cyc15 (purple) tended to have small to medium sizes, and were located near the border between larger cells and smaller cells. The median features displayed by each putatively identified neuron (Fig. 9B) suggest that most neurons do not possess an individual feature that uniquely identifies them in this feature-space. Rather, most neurons display a unique combination of many features. For example, even though neurons cyc10 and cyc11 are small cells among many other small neurons (e.g., ganglia d–g in Fig. 9A), and share many features they can often be identified due to their unique combinations of features (Fig. 9B). This also appears to be the case for many neurons from the rostral surface (Fig. 10A, B), where most putatively identified cells were small neurons that were responsive to stimulation of many nerves. Similar to caudal neurons, patterns were clear in many putatively identified rostral neurons, such as the sizes and positions of cyc03, cyc04, cyc06, cyc14, and cyc16 (Fig. 10). Taken together, these findings suggest that unsupervised functional neurocartography of the *Aplysia* buccal ganglion can overcome intersubject variability and consistently identify putatively homologous units across animals.

**Figure 9.**
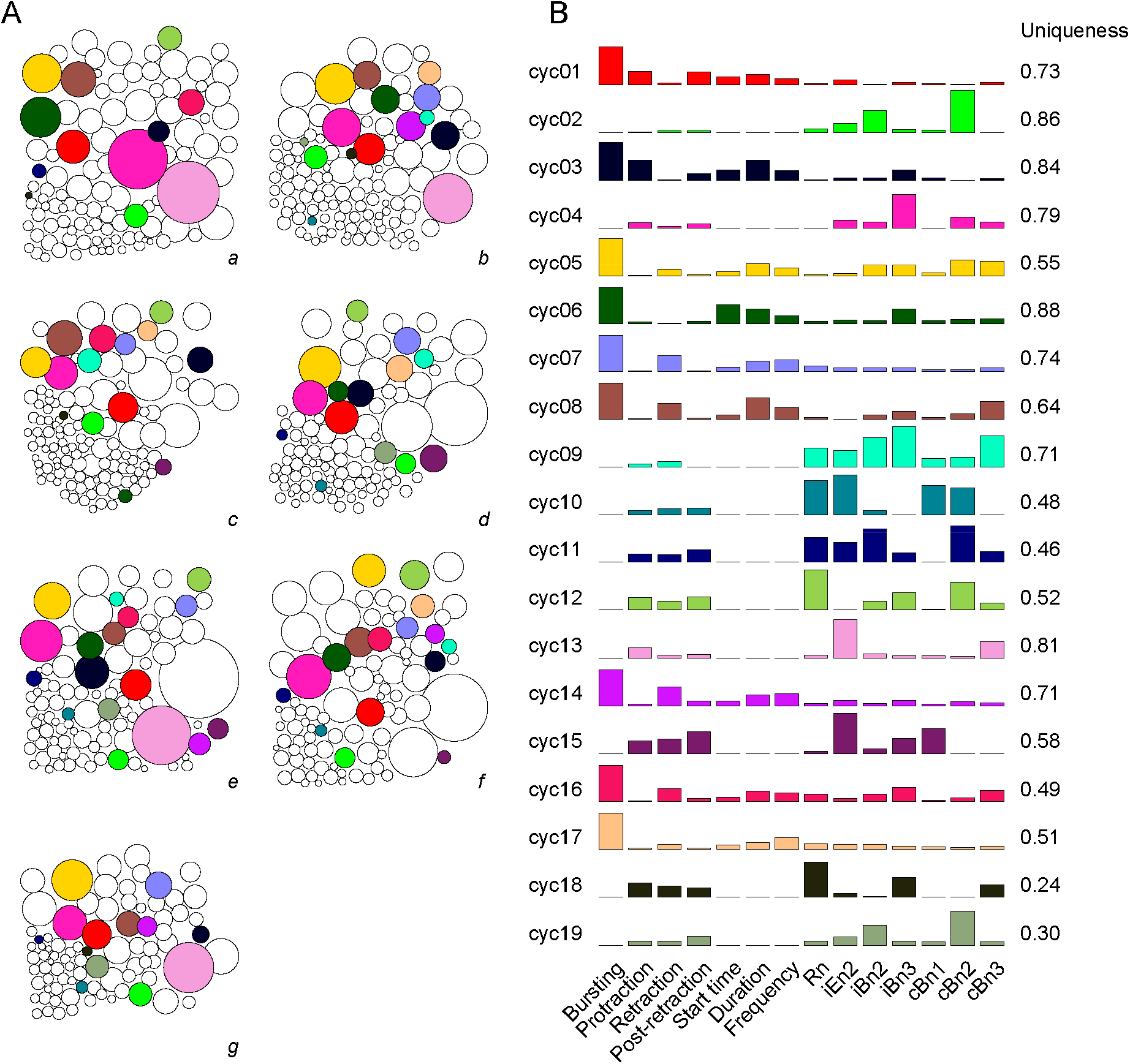
Overview of putatively homologous neurons across seven caudal preparations. **A.** Diagrams representing the sizes and positions of individual neurons recorded in each caudal preparation (*a–g*). Filled circles indicate putatively homologous neurons, with each color denoting one individual group. For example, the yellow circle (cyc05) usually located on the upper left quadrant of each preparation indicates a neuron that was putatively identified in all seven ganglia. Note that, although many neurons exhibit roughly consistent localization across ganglia, there is substantial variability in the exact position of each cell. **B.** Overall feature profiles of each group. For each feature, the median value across all members of a group is displayed. The y-axis for each feature (i.e., column) is normalized to the maximum observed value of that feature across all caudal neurons. Color scheme corresponds to neurons in panel A. Group labels (cyc01–19) correspond to those in Fig. 8A.

**Figure 10.**
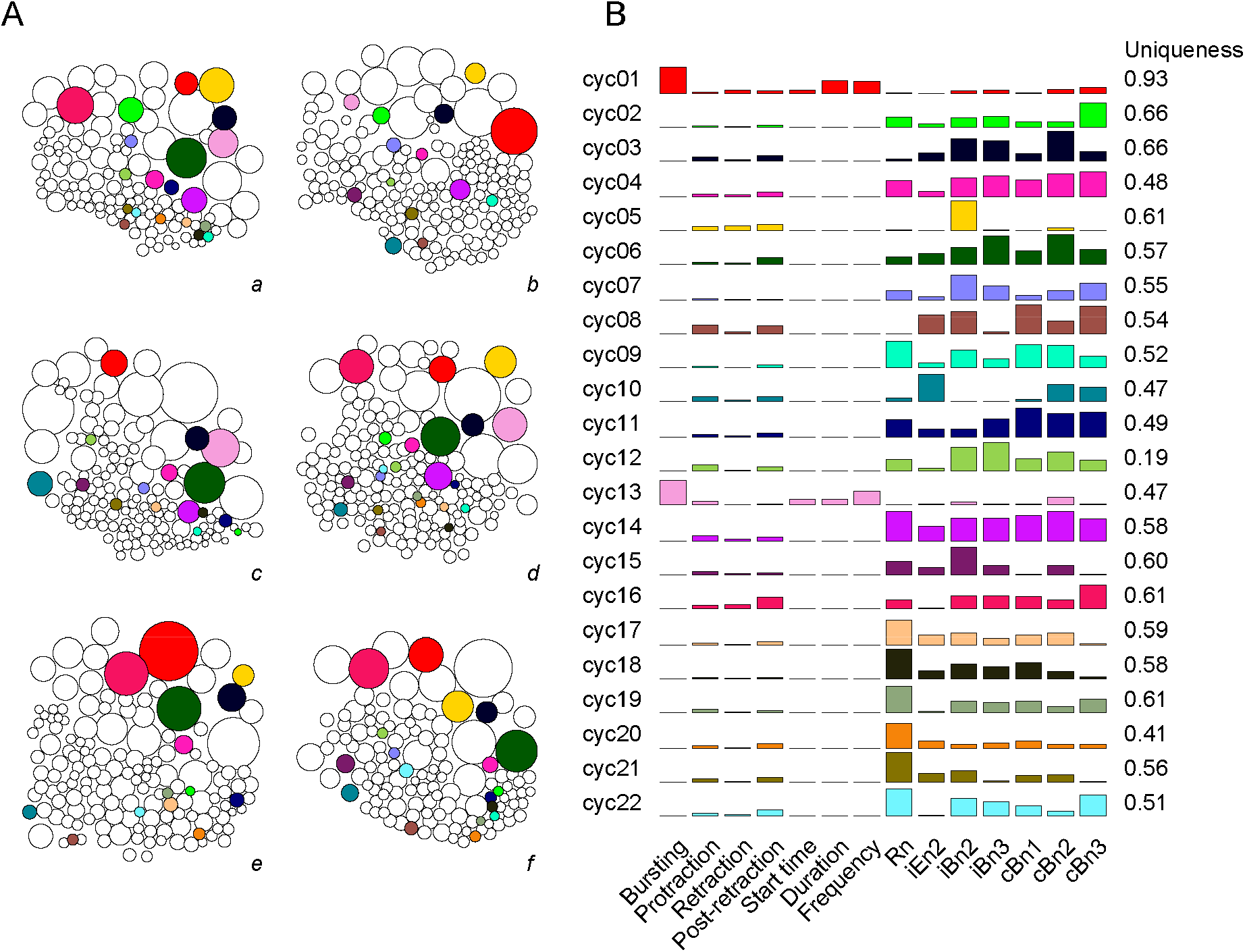
Overview of putatively homologous neurons across six rostral preparations. Same as Figure 9, but for rostral preparations. **A.** Diagrams representing the sizes and positions of individual neurons recorded in each rostral preparation (*a–f*). **B.** Overall feature profiles of each group.

## DISCUSSION

### *Functional neurocartography of the* Aplysia *buccal ganglion*

The present study applied the functional neurocartography framework (Frady et al., 2016) to the buccal ganglion of *Aplysia* and made several key modifications to enhance its scalability. Neurocartography is composed of two central parts: i) construction of the feature-space, and ii) intersubject matching of neurons based on that feature-space. Each of these is discussed in the following paragraphs.

### Developing the feature space for the buccal ganglion

The original framework of neurocartography was developed using VSD imaging data from the leech ganglion. Even though the framework itself is highly versatile, many functional features are specific to each system. Therefore, the first step of our analysis was establishing the feature-space for the buccal ganglion that contained the optimal set of features. The automated algorithm selected parameters related to BMPs. This is not surprising given that many neurons in the buccal ganglion have been characterized based on their activity during BMPs (Baxter and Byrne, 2006; Cropper et al., 2004; Elliot and Susswein, 2002; Jing and Weiss, 2005; Morton and Chiel, 1993b). In addition to purely functional features, the VSD signals evoked in response to the stimulation of different nerves (Fig. 3) were used, which aided discrimination of neurons. Feature selection based on within-subject variability (Fig. 4) demonstrated that stimulation of various nerves provided useful information in both caudal and rostral datasets. In contrast, some features that were useful for the caudal surface were not useful for the rostral dataset (e.g., protraction and retraction). The rejection of protraction and retraction in the rostral dataset was surprising given that the phasing of neurons has historically been an important defining feature for neurons in the buccal ganglia. One possible explanation is that, for the purpose of large-scale neuron discrimination, the information these features bore was redundant with information provided by other features. In that case, it would be beneficial for the overall matching to discard such a redundant feature, even when it is both information-rich and salient to experimenters. These results suggest the need to carefully identify and evaluate the feature sets that are optimally informative for each neuronal population of interest.

### Unsupervised pipeline for matching of putatively homologous neurons across subjects

One of our primary goals was to increase the degree of automation, thereby enhancing scalability. The original framework (Frady et al., 2016) was designed to utilize human instruction to manage intersubject variability through skewing of the feature-space. In addition, it relied on manual processing to “stitch” the collection of pairwise matching results into groups of putatively homologous neurons. Importantly, this process is nontrivial because pairwise matches are not always consistent if more than two animals are considered.

A key tool in our approach was cross-animal matching based on cyclic matching (Hofbauer, 2016). Although cyclic matching can avoid the problem of inconsistency across pairwise matches, its matching results are generally sensitive to the order of experiments (subjects). Notwithstanding, cyclic matching can be run with many randomly permuted orders of experiments. This strategy makes it possible to discriminate between neuron pairs that are consistently matched across permutations and those that are not (copresence analysis; Fig. 6B).

The parametrization of individual neuron pairs through the copresence analysis opened a path to the automation of feature-skewing (Fig. 7), allowing for the entire analytic pipeline, including matching of putatively homologous neurons, to be automated and unsupervised. Thus, the combination of cyclic matching with the copresence analysis is a promising scalable approach for functional neurocartography. It should be noted, however, that this approach would not work well when the number of subjects is small because of the limitation in the number of possible random permutations. In principle, the ability to estimate intersubject variability is dependent upon the number of subjects from which the estimation is performed. Thus, it is to be expected that a reduced number of animals should impair performance of any approach that needs to estimate intersubject variability. It is worthwhile to point out that the original manual framework is well suited for cases where the number of subjects is prohibitively small for the automated approach.

### Limitations and further improvements

Several aspects of the approach may be improved in future studies. First, inclusion of additional informative features would increase the number of identifiable neurons. Many neurons were not responsive to nerve stimulation, did not display characteristic activity during BMPs, and often had similar spatial features; thus, there was little information available to differentiate these cells. Therefore, one direct means to increase the number of identifiable homologous neurons would be to uncover additional features that are particularly informative for this population. Possible features to be examined include: i) response to stimulation of other nerves or connectives (e.g., i/cEn1, cEn2, i/c salivary nerves, i/c I2 nerves, cRn, iBn1, i/c cerebral-buccal connectives); ii) activity responses to various chemical manipulations (e.g., ionic concentrations of the artificial seawater bath solution, small molecule neurotransmitters, neuropeptides, receptor agonists and/or antagonists); iii) immunostaining for specific molecules (e.g., neurotransmitters, neuropeptides, receptors, kinases, etc.); and iv) patterns of synaptic connectivity.

Second, a reduction in the sparseness of the current features could be beneficial. The feature set in this study includes a larger number of features than Frady et al. (2016) and, more importantly, those individual features appeared to be sparser. This may have made the matching analysis more prone to measurement noise and compromised the accuracy of matching, especially in neurons with relatively few prominent features. To further improve matching accuracy by increasing robustness to noise, the most straightforward approach would be to examine whether simple dimensionality reduction techniques (e.g., principal component analysis) would effectively reduce the number of features without losing critical information, thereby reducing feature sparseness.

Third, estimates of intersubject variability may be useful for feature selection. In both the original framework and in the present study, feature selection was based on the within-subject variability of feature data. This is purely due to the lack of tools to evaluate features based on their intersubject variability, as intersubject variability is more directly relevant to intersubject matching of neurons. The automated matching procedure developed in this study could allow for the optimization of features based on their intersubject variability if various subsets of features were directly fed to the matching pipeline. Although the metrics for evaluating the overall performance of matching results are still limited, this strategy could simplify and improve the process of feature selection.

Finally, the analytical pipeline may be further refined through its application to a dataset for which the ground-truth is known. We had limited means to assess the accuracy of matching results, primarily due to the lack of ground-truth data. Although there is no experimental technique existing to generate such large-scale ground-truth data, the use of noise-added versions of real data may be helpful in assessing the performance of the analytical pipeline. For example, Yu et al. (2021) successfully used semi-synthetic data to train an artificial neural network to match neurons across subjects in *C. elegans*. Furthermore, experiments could be designed to validate a small subset of the results by performing intracellular recordings following the imaging, similar to Frady et al. (2016), given that many neurons in the buccal ganglion have been electrophysiologically and morphologically identified (e.g., Church et al., 1991; Dembrow et al., 2003; Gardner, 1971; Kabotyanski et al., 1998; Plummer and Kirk, 1990; Susswein and Byrne, 1988). An ultimate goal in the neurocartography of the *Aplysia* buccal ganglion would be to associate the groups of homologous neurons with neurons that were previously characterized in the literature. Such an effort may lead to groups of homologous neurons with no previously identified counterpart, i.e., groups of yet uncharacterized neurons, and ultimately to a thorough canonical map of the *Aplysia* buccal ganglion.

## MATERIALS AND METHODS

Experimental methods in this study closely follow those described in Costa et al. (2022).

### Animals and preparations

*Aplysia californica* were obtained from South Coast Bio-Marine LLC (San Pedro, CA) or bred at the National Resource for *Aplysia* of the University of Miami (Miami, FL). Data were collected from a total of 13 animals weighing between 25 and 50 g. Prior to dissection, animals were injected with a volume of isotonic MgCl_2_ equal in milliliters to 50% of the animals’ mass in grams. The buccal mass with buccal ganglia attached was isolated from the animal and placed in a Sylgard-coated chamber containing artificial seawater with a high concentration of divalent cations (HiDi-ASW; 210 mM NaCl, 10 mM KCl, 33 mM CaCl_2_, 145 mM MgCl_2_, 20 mM MgSO_4_, 10 mM HEPES, pH adjusted to 7.5 with NaOH). HiDi-ASW was used to reduce activity and muscle contractions during dissection. Buccal ganglia with buccal nerves were excised from the buccal mass and pinned down. The sheath of connective tissue around the ganglia was kept intact. Bipolar suction electrodes for recording and stimulation were positioned on each nerve (see below). HiDi-ASW was substituted with normal artificial seawater (ASW; 450 mM NaCl, 10 mM KCl, 10 mM CaCl_2_, 30 mM MgCl_2_, 20 mM MgSO_4_, 10 mM HEPES, pH adjusted to 7.5 with NaOH) prior to recording. Preparations were maintained at 15 °C throughout the experiment.

### Extracellular electrophysiology

Bipolar suction electrodes were used to monitor nerve activity associated with buccal motor patterns (BMPs). Nerve signals were amplified by A-M Systems model 1700 differential AC amplifiers and digitized by Axon Instruments Digidata 1322A at 20 kHz. The following nerves were recorded: the ipsilateral and contralateral buccal nerves 2 and 3 (iBn2, cBn2, iBn3, cBn3); contralateral buccal nerve 1 (cBn1); ipsilateral radula nerve (iRn); and ipsilateral esophageal nerve 2 (iEn2).

The responses of the neurons in the imaged hemiganglion to brief electric shocks to the peripheral nerves of the ganglion were characterized. This procedure was designed to activate neurons directly if a particular neuron had an axonal projection in the stimulated nerve, or indirectly via synaptic drive from other neurons, or a combination of the two. This procedure was repeated over 40 trials for each of the nerves (i.e., iBn2, iBn3, iEn2, iRn, cBn1, cBn2, cBn3). Stimulation was automatically triggered by a WPI Pulsemaster A300 and delivered by a WPI 850A stimulus isolator. Stimulus intensity, pulse duration and frequency were respectively 15 V, 0.5 ms, 2 Hz for caudal and 30 V, 1 ms, 2 Hz for rostral preparations. Both types of stimulation were effective in eliciting responses and produced informative features.

BMPs were manually identified from nerve activity. Activity in Bn1 was used as an indicator of the timing of each BMP phase. Bn1 is particularly informative because it is active during protraction, inactive during retraction, and active again upon retraction termination. A BMP was identified when three criteria were met: (1) large-unit activity lasting at least 3 s occurred in Bn1; (2) the Bn1 activity was followed by large-unit activity in at least one of the two Bn3s; (3) large-unit Rn activity occurred during either the Bn1 activity (protraction) or the subsequent Bn1 suppression (retraction). Large-unit activity was defined as all activity with amplitude above a voltage threshold for each nerve, which was set independently for each experiment. The threshold for Bn1 was set to include all units that were consistently active during protraction and inactive during retraction. The Bn3 threshold was set to include only the largest units, which correspond to B4, a neuron known to be active at the onset of retraction. Large-unit activity was counted when at least 3 spikes occurred with interspike intervals ≤4 s.

### Voltage-sensitive dye (VSD) imaging

Isolated buccal ganglia were stained for 7 min with 0.2 mg/ml of the absorbance VSD RH-155 (TRC Canada) in ASW. After staining, the solution was swapped for a lower concentration of RH-155 in ASW (0.01 mg/ml), which was kept for the remainder of the experiment. Using a Gilson Minipuls 3 peristaltic pump, the solution was continuously cycled through a Warner Instruments SC-20 inline cooler/heater under control of a Warner Instruments CL-100 temperature controller, which maintained the preparation at 15 °C. Imaging data was acquired by a Deep Well NeuroCMOS 128×128 camera (RedShirtImaging, SciMeasure) sampled at 1 kHz. The camera was fitted to an Olympus BX50WI microscope equipped with an Olympus 20× 0.95 NA XLUMPlanFL water immersion objective lens. The size of the field of view was 820×820 μm. The preparation was illuminated with a 150 W, 24 V Osram halogen light bulb powered by a Kepco ATE 75-8M power supply. Before reaching the preparation, light passed through a Brightline 710/40 nm bandpass filter.

Preparations were imaged during BMP generation for two separate 120 s time windows. For one rostral preparation, no BMPs were captured during one of the time windows. Thus, that imaging time window was repeated. After the second window, preparations were imaged during stimulation of peripheral nerves (20 s per nerve; see *Extracellular electrophysiology* above). The total imaging time per preparation was approximately 380 s. Either the rostral or caudal surface of the left buccal hemiganglion was imaged. The field of view was adjusted so that the center portion of the hemiganglion was visible. Approximately 80% of the total surface area was visible on each preparation. The focal plane was set to the region with the greatest number of neurons. Neurons in the ventral cluster, as well as B1 and B2 when imaging the caudal surface, were consistently in the field of view. However, neurons close to the edges of the field of view may not have been captured in all preparations (e.g., B4/5, B8a/b, dorsal portions of the sensory cluster). The resulting position of the ganglion was reasonably consistent across experiments. In addition, variations were accounted for during data analysis by approximating the orientation and offset of both the ventral cluster and the entire ganglion, and then aligning each recording accordingly, as previously described (Neveu et al., 2017).

### Data processing

VSD data were processed in a similar manner to that described elsewhere (Neveu et al., 2017; Costa et al., 2022). Briefly, regions of interest (ROIs) were drawn around the soma of each neuron, and the light intensity was averaged across pixels within the region for each time point. Traces for each cell were taken as the change in light intensity relative to baseline (i.e., ΔI/I), where baseline was defined as an average of the first 10 frames after shutter opening. Traces were passed through a bandpass butterworth filter (Fpass1 = 15 Hz, Fstop1 = 0.1 Hz, Fpass2 = 140 Hz, Fstop2 = 0.5 kHz, Apass = 0.1, Astop1 = 60, Astop2 = 60).

Similar to Costa et al. (2022), principal component analysis (PCA) was used to isolate and remove correlated noise, such as that associated with sample movement. PCA extracts a set of components from the data that can be ordered by the amount of variance-covariance in the data explained by each component. If some components consist primarily of noise, the data can be reconstructed without those components to improve signal-to-noise ratio. At the millisecond timescale, most of the correlation between the traces of individual neurons was due to noise that affected many neurons simultaneously, such as tissue movement, vibrations, and fluctuations in illumination. This was the case because the amplitude of RH-155 signals is very small (i.e., in the order of 1×10^-4^ ΔI/I), and, importantly, because action potentials in *Aplysia* are not correlated at the millisecond scale during spontaneous BMP generation. Thus, PCA was performed on the traces from all neurons in a preparation, removed the top n components accounting for 85% of the variation, and reconstructed the data. Upon visual inspection of signal quality this threshold consistently reduced noise without signal decrement.

Spike detection was performed by the algorithm described in Neveu et al. (2017). Briefly, spikes were detected in a neuron when a downward deflection in the VSD trace exceeded 2.75 times the standard deviation of the trace and the peak of this downward deflection was followed after 7 ms by an upward deflection in excess of 3.25 times the standard deviation. The re-arm window for spike detection was set to 22.5 ms.

### Feature extraction

*Position* and *size* of a neuron were determined based on the hand-drawn ROIs overlying individual neurons in a pixel coordinate of the original data (128×128 pixels). The position of a neuron was defined as the *x* and *y* coordinates of the centroid of the pixels included in the ROI. The size was defined as the total number of pixels of the ROI. A small subset of neurons was located on the edge of the camera sensor, and thus a part of their cell bodies was not within the field of view. No adjustment to size or position was made to account for this.

All feature parameters determined from the VSD imaging of BMP activity were based on the spike timing data obtained from the basic data processing described in the preceding subsection. The data from two imaging windows (see *Voltage-sensitive dye (VSD) imaging*) were concatenated. *Phase preference* features (*protraction*, *retraction*, and *post-retraction*) were defined based on a simple firing rate during each of the BMP phases. The number of spikes during the phase of interest and the duration of phase were summed across all the occurrences of BMPs. The ratio of the summed spike counts over the summed duration was considered the firing rate. *Phase preference* is defined as a relative firing rate that is normalized to the highest firing rate across all neurons for a given BMP phase of a given animal.

The properties of burst-like activity were used to obtain additional features. For this purpose, the burst detection algorithm described in Neveu et al. (2017) was used. Briefly, bursts were identified whenever 3 spikes occurred with ≤400 ms interspike interval, and when the total firing rate within bursts was significantly different from the firing rate outside of bursts. Neurons were assigned true for the binary feature *bursting* if they displayed burst-like activity, and false otherwise. *Frequency* was defined as the overall firing rate within burst occurring during BMPs for a given neuron, regardless of phase. The temporal properties of burst-like activities were measured after normalizing all time points during a given BMP phase to the duration of that phase. This approach allowed for the relative timing of bursts to be captured, regardless of variability in the duration of phases and BMPs. *Start time* was defined as the time elapsed between BMP onset and onset of the first burst for each neuron. *Duration* was defined as the sum of the durations of all bursts occurring during BMPs for a given neuron. When the *bursting* feature was false, the *frequency*, *start time*, and *duration* features of that neuron were set to zero.

Feature parameters from the nerve stimulation experiments were based on normalized kernel time courses (ΔI/I) that were not temporally filtered or denoised because processing would significantly alter the waveform of response. At first, a stimulus-triggered average of response was calculated for each combination of neuron and stimulated nerve. The VSD time course was super-sampled 10 times by replacing each data point with 10 data points of the identical value. The time points of electrical shock were extracted from the nerve recording and corresponding time points were found in the super-sampled VSD time course. VSD data of a time-window from −100 ms to +500 ms of stimulus was extracted for each occurrence of electrical stimulus and then their mean time courses were calculated. In each averaged time course, constant offset of the baseline was corrected by subtracting the mean across a time window from −20 to 0 ms from all data points. The amplitude of response was defined as the integral of -ΔI/I in a time window from 0 to 25 ms. Although this time window was potentially too short for a subset of long-lasting responses, the contribution of baseline drift became detrimental when longer time windows were considered.

The feature parameters were also calculated from the half-split data to assess their discriminability (see the next subsection). Data from spontaneous activity recordings were split into the odd number and even number occurrences of BMPs after pooling the two imaging windows. Nerve-stimulation data were split into the odd and even number trials of electrical stimulation. No half-split data were considered for the position and size parameters as they were unchanging throughout the experiment.

### Feature discriminability

The calculation of the discriminability index was done by dividing the data within a single experiment into two separate halves and comparing the neuron in one half to itself, as well as to the most similar non-self neuron, in the other half. The comparisons were performed by applying the whitening transform to the entire feature-space and then considering each feature as an independent dimension in an *F*-dimensional space. Each neuron, then, could be represented as a vector in this space and comparing two neurons could be done by computing the *ℓ*_2_ norm of the vector distance between them. Thus, the *discriminability* of a given neuron *i* in feature-space *f* was given by the same equation used in Frady et al. (2016):

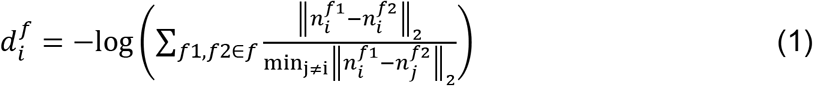

where *f1* and *f2* represent two halves within the same subset of features represented by *f*. The parenthetical expression represents the comparison of a neuron’s self-similarity between the two halves to its most similar non-self neighbor in the feature-space. By taking the negative logarithm of this fraction, positive values are assigned to self-similar neurons and negative values to non-self-similar ones. For a neuron *i*, if this value was positive, the neuron was said to be *discriminable* in this feature-space.

The subset of features that was most beneficial for identifying neurons across experiments should maximize the number of discriminable neurons. The total *feature-space discriminability* was given by the following expression:

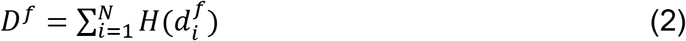

where *H(x)* represents the Heaviside step function, used to convert positive discriminabilities into ones, which could then be tallied up to represent the total discriminability of feature-space *f*. Consequently, the feature-space is maximally informative when *D* is maximized.

To determine the optimal subset of features to maximize the feature-space discriminability, at first, a subtractive assessment of discriminability was employed: starting with the full feature set, each feature was removed one-by-one and discriminability was reassessed. If discriminability decreased with the removal, the feature was considered informative and kept in the feature set. Otherwise, the feature was considered not helpful and removed. Next, because how informative a feature is may be dependent on the redundancy of information with other features, the once-removed features were added back one by one, and discriminability was reassessed. If the added feature increased discriminability, it was retained in the feature space.

Compared to the use of the discriminability index by Frady et al. (2016), there was a departure in the present study. Only functional features, but not spatial ones, were used for assessment because half-split data are unavailable for the spatial features. Although previously this was addressed by intentionally adding noise to simulate biological variation (Frady et al., 2016), the amount of added noise directly impacts the discriminability score. Therefore, in this study, the analysis of discriminability was rather focused on the functional features with the premise that spatial features are informative.

### Pairwise matching algorithm

The pairwise matching of neurons between experiments were based on the feature-space optimized for the highest discriminability. First, the whitening transform was applied to the entire feature-space for all experiments simultaneously. This transform set all the feature-space covariance matrix to the identity matrix, and normalized the values of the features, which had varying magnitudes and scales depending on the respective features.

Each neuron in one experiment was compared to all neurons in another by considering the distance of feature vectors between them. The vector distance was then converted to the matching strength, which takes into account the uniqueness of match and is defined as follows:

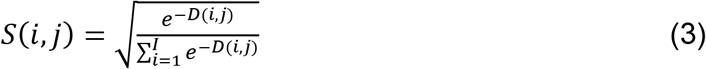

where *I* represents the set of neurons in a given experiment being compared to neuron *j* in the other and neuron *i* is the neuron of interest. This expression, therefore, ranges from 0, where the distance *D* between the two neurons of interest is infinite, to 1, where *D* is 0 and all other distances to neuron *j* are infinite, meaning neuron *i* is uniquely and strongly matched to neuron *j*.

Matching pairs were determined through the Kuhn-Munkres assignment algorithm (Kuhn, 1955; also referred to as the *Hungarian method*), which enabled the combinatorial optimization, maximizing the overall strength of matches while accepting the presence of pairs with non-highest match strength. The neuron-to-neuron matches were further optimized for intersubject variability through a skewing of the feature-space of interest, where the magnitude of the distance vectors between these neurons was calculated similarly to before; although the vectors representing each neuron now had each a weight matrix assigned to it, as given by the following:

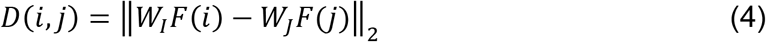

As illustrated by the equation, the weight matrices are defined per experiment. Weight matrices *W_I_* and *W_J_* were determined with the algorithm implemented in the neurocartography software (see *Feature skewing*).

### Cross-animal evaluation of pairwise matching

The results of pairwise matching were further analyzed regarding their consistency across multiple animals. Specifically, groups of neurons satisfying the following criterion were sought: every member in a group of four neurons was matched to all three other members of the group by pairwise matching. Then, separate groups of four neurons that shared member neurons were merged. This merging tolerated a subset of neuron pairs in the final merged group not being matched in pairwise matching.

For a given neuron in the entire dataset (seed neuron), all its pairwise-matched neurons were identified (from 6 other subjects in the caudal dataset or 5 subjects in the rostral dataset, except for missing matches due to unequal neuron numbers between subjects). Next, for a given pair of the seed neuron and its directly-matched neuron, their pairwise counterparts in the remaining subjects were compared and kept only if they were the same. If more than two neurons were kept as shared counterparts of the seed and its direct-match, every possible pair among the shared counterparts was listed and tested in pairwise matching. If matched, the seed neuron, the seed’s direct-match, and the shared counterpart pair were saved as a perfect group of four, in which every possible pair is a best match in pairwise comparisons. This procedure was repeated taking every neuron in the dataset as the seed neuron. From the resulting list of perfect groups of four, redundant entries were removed, then entries sharing the same neuron(s) were merged in a recursive manner. In case a merged group contained two neurons from the same subject, both of them were removed. The final results (i.e., a list of successful groups) were manually inspected for additional irregularities (none was detected).

### Cyclic matching

As part of pairwise matching, the neuron-to-neuron match strength was computed between all combinations of neurons for all possible pairs of experiments. For a given neuron in one experiment, every neuron from the experiments to be compared were ranked based on their match strengths. Such preference ranks were determined for every neuron in the dataset for every possible experiment to be paired.

Families of neurons with cyclic matching preferences were identified based on Hofbauer (2016) (Fig. 6A). Briefly, the order of experiments in the matching cycle was defined at first by random assignment. Starting with the first experiment in the cyclic order, the neurons in this first experiment were each matched to neurons in a second experiment as in the pairwise matching (see above). These newly added neurons were then matched to neurons in the next experiment, until the last experiment. For neurons in the last experiment, the neurons to be matched in the first experiment had been already determined (i.e., the first neuron in each family), although these final matches do not necessarily coincide with matches that would result from pairwise matching. Therefore, the pairwise matching strength was used to determine the preference rank of the first neuron in each family with respect to the last (preceding paragraph). The families of matches that had the best last-to-first preference rank were considered most content (highest contentness) and removed from the pool. Then the preference ranks of the neurons in the remaining matching series were updated and the same process was performed iteratively, until no more neurons remained in the experiment with the fewest neurons.

For the copresence analysis (Figure 6B), the cyclic matching described above was repeated 100 times with different orders of experiments, which were randomly permuted with no identical orders allowed. One neuron in the dataset was picked as the reference neuron, and for every neuron from every experiment except the one that includes the reference neuron, the number of runs in which the neuron of interest and the reference neuron were placed in the same family was counted and termed copresence count. Because the number of neurons in an experiment can be larger than the number of families, it is possible that the reference neuron is not placed in any of the families. The number of runs in which the reference neuron is placed in any of the families is termed appearance count, and this value sets the upper boundary of copresence count for a given reference neuron. The analysis of copresence count was performed taking every neuron in the dataset as the reference neuron.

### Feature skewing

Skewing of the feature-space was performed with a custom algorithm implemented in the Imaging Computational Microscope (ICM; Frady and Kristan, 2015). Briefly, the algorithm, termed weighted correspondence minimization (WCM), finds a transform of the feature-space that minimizes the distance among a set of selected matches between neurons. The WCM algorithm was not modified. Instead, an approach was developed to automate selection of matches to be optimized by the WCM transform. This approach was based on the copresence profiles obtained from running multiple permutations of the cyclic matching algorithm (described in the previous section). When a target neuron’s copresence count proportion relative to the appearance count of a reference neuron surpassed a given threshold, the pair of neurons was taken as selected matches. The total set of selected matches was used as input to the WCM algorithm, which, therefore, sought a transform of the feature-space that minimized the distances among these selected matches. This transform corresponds to the weight matrices used to compute vector distances between neurons following skewing of the feature-space (i.e., *W_I_* and *W_J_* in equation 4).

The quality of the resulting feature-space was assessed by the sparseness metric

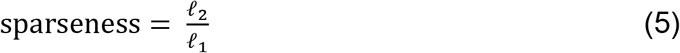

where *ℓ*_p_ is the p-norm of the copresence profile vector. A sparse vector has only a few non-zero dimensions, whereas a non-sparse vector has many non-zero dimensions. Thus, both the *ℓ*_1_ norm (i.e., the modulus) and the *ℓ*_2_ norm (i.e., the Euclidean distance from the origin) increase as sparseness decreases, but the *ℓ*_1_ norm does so at a faster rate. However, either norm has the disadvantage of being scale-dependent. In the particular case of the copresence profile, reducing the appearances of a reference neuron would lead to lower *ℓ*_1_ and *ℓ*_2_ norms of the copresence vector, implying an increase in sparseness without a clear improvement in performance. Conversely, the ratio between the norms offer a scale-invariant alternative (Krishnan et al., 2011; Yin et al., 2014). The ratio will only imply an increase in sparseness when the distribution of copresence counts for a given reference neuron converges on fewer partner neurons, which indicates that matches are more consistent and less ambiguous across runs. Thus, this sparseness metric was used to compare the unskewed feature-space with the skewed feature-spaces using a range of copresence proportion thresholds (Fig. 7).

### Identification and characterization of canonical neurons

Based on the copresence counts, putatively homologous neurons were grouped using the following procedure. To obtain the copresence counts used in this process, 100 runs of cyclic matching were repeated 10 times, and mean copresence counts were calculated across the repeats for each pair of the reference neurons and the neuron of interest. Then, for a given reference neuron N_1_ and neurons in the experiment under comparison, the neuron with the highest copresence count was identified as N_2_ and its uniqueness index (UI) was defined as follows:

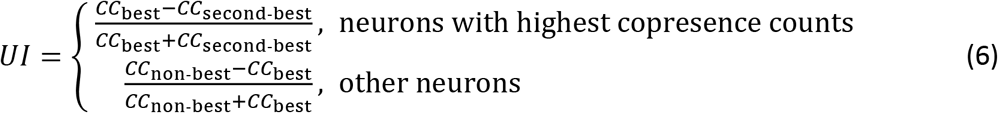

where CC_best_ is the copresence count of N_2_ and CC_second-best_ is that of the neuron with the second-highest count of the same experiment after N_2_. (The definition of UI is extended for neurons that do not have the highest count and by the second equation, where CC_non-best_ is the copresence count of that neuron.) When the UI was higher than 0.5, N_2_ was set as the new reference neuron and it was tested whether N_1_ showed the highest copresence count in the experiment under comparison with UI above 0.5. If true, N_1_ and N_2_ were considered as an established pair of neurons. From the UI equation it follows that a value of 0.5 is equivalent to a copresence count that is three times higher for the best neuron than the count of the second-best. This value was chosen as a threshold to have minimal false positives considering the standard error of the mean copresence counts. Next, for each neuron that formed an established pair, all established pairs including that neuron were gathered as a preliminary group. For the preliminary groups, if one is identical to or a subset of another preliminary group, they were integrated into a single preliminary group. In case a single preliminary group contained two member neurons from the same experiment at this point, both of them were removed. Finally, if two preliminary groups shared a subset of members but each also had its unique member(s) (e.g., U_1_ from one preliminary group and U_2_ from another, it was tested whether U_1_ made an established match with a different neuron from the experiment including U_2_, or vice versa). If neither is true (i.e., there was no explicit counterevidence), they were allowed to be merged as a single group. If either (or both) is true, U_1_ and U_2_ were removed and only shared members were kept in the merged preliminary group. Preliminary groups that had four or more member neurons at this point were defined as the groups of putatively homologous neurons.

For each group of putatively homologous neurons based on cyclic matching, all unskewed feature parameters of member neurons were retrieved and the median across member neurons was calculated for each feature. The resulting set of median feature parameters were considered as the features representing the group of homologous neurons (i.e., a canonical neuron).

### Code and data availability

Code and data will be deposited to public repositories following peer-review.

## Acknowledgments

We thank Dr. E. Paxon Frady for assistance with the implementation of the original functional neurocartography framework. This work was supported by National Institutes of Health grant R01 NS101356 (JHB), CNPq Science without Borders Scholarship 203059/2014-0 (RMC), and a Larry Deaven Ph.D. Fellowship in Biomedical Sciences (RMC).

